# Chlorophyll fluorescence-based estimates of photosynthetic electron transport in Arctic phytoplankton assemblages

**DOI:** 10.1101/2021.08.17.456659

**Authors:** Yayla Sezginer, David J. Suggett, Robert W. Izett, Philippe D. Tortell

## Abstract

We employed Fast Repetition Rate fluorometry for high-resolution mapping of marine phytoplankton photophysiology and primary productivity in the Lancaster Sound and Barrow Strait regions of the Canadian Arctic Archipelago in the summer of 2019. Continuous ship-board analysis of chlorophyll *a* variable fluorescence demonstrated relatively low photochemical efficiency over most of the cruise-track, with the exception of localized regions within Barrow Strait where there was increased vertical mixing and proximity to land-based nutrient sources. Along the full transect, we observed strong non-photochemical quenching of chlorophyll fluorescence, with relaxation times longer than the 5-minute period used for dark acclimation. Such long-term quenching effects complicate continuous underway acquisition of fluorescence amplitude-based estimates of photosynthetic electron transport rates, which rely on dark acclimation of samples. As an alternative, we employed a new algorithm to derive electron transport rates based on analysis of fluorescence relaxation kinetics, which does not require dark acclimation. Direct comparison of kinetics- and amplitude-based electron transport rate measurements demonstrated kinetic-based estimates were, on average, 2-fold higher than amplitude-based values. The magnitude of decoupling between the two electron transport rate estimates increased in association with photophysiological diagnostics of nutrient stress. Discrepancies between electron transport rate estimates likely resulted from the use of different photophysiological parameters to derive the kinetics- and amplitude-based algorithms, and choice of numerical model used to fit variable fluorescence curves and analyze fluorescence kinetics under actinic light. Our results highlight environmental and methodological influences on fluorescence-based productivity estimates, and prompt discussion of best-practices for future underway fluorescence-based efforts to monitor phytoplankton photosynthesis.

## Introduction

Phytoplankton productivity in polar marine waters is constrained by nutrient and light availability, which fluctuate dramatically across seasonal cycles and shorter time and space scales [1]. In late summer, when sea ice cover is at a minimum and the mixed layer is shallow and highly stratified, phytoplankton are exposed to high solar irradiances and low nutrient concentrations [2]. Under these conditions, the growth and photosynthetic efficiency of Arctic phytoplankton becomes nitrogen-limited [3]–[5]. However, localized regions of elevated productivity can persist where various processes transport nutrients into the mixed layer, including upwelling, tidal mixing, and freshwater input from rivers and glaciers [6]–[8]. Modelling studies suggest that Barrow Strait in the Canadian Arctic Archipelago is one such productivity hotspot, with strong tidal currents and shallow sills driving vertical mixing in a region where Pacific and Atlantic-derived water masses converge [9], [10]. Additionally, Barrow Strait receives glacial and land-derived nutrients from the Cornwallis and Devon Island rivers [11], [12]. Rapid climate change in the Arctic is expected to have complex effects on these nutrient delivery mechanisms through the intensification of coastal erosion [13], increasing river inputs [14] and reduced vertical mixing due to intensifying stratification [15], [16]. At present, it is unclear how phytoplankton productivity will respond to these anticipated perturbations. Assessing phytoplankton productivity in physically-dynamic marine waters requires high spatial resolution measurements that cannot be obtained from traditional discrete bottle incubation methods, such as ^14^C uptake experiments. For this reason, oceanographic field studies have increasingly employed continuous sampling of surface water properties using variable chlorophyll *a* (Chl*a*) fluorescence from Fast Repetition Rate Fluorometry (FRRf) and other related methods to rapidly and autonomously assess phytoplankton photophysiology and productivity (e.g. [17]–[21]). Such variable fluorescence techniques rely on the inverse relationship between Chl*a* fluorescence and photochemistry. These processes, along with heat dissipation, represent the three energy dissipation pathways of absorbed light energy within Photosystem II (PSII) [22]. Fast Repetition Rate Fluorometry operates by supplying rapid excitation light pulses to progressively saturate the photosynthetic pathway and simultaneously induce a measurable Chl*a* fluorescence response – often referred to as a fluorescence transient [23]. Analysis of Chl*a* fluorescence transients provides information on the photochemical efficiency and functional absorption cross section of PSII, as well as estimates of the turnover rate of photosynthetic electron transport chain molecules [24], [25]. Fast Repetition Rate fluorescence transients can be obtained nearly instantaneously, offering opportunities for very high temporal and spatial resolution measurements.

Primary productivity is typically estimated from variable Chl*a* fluorescence measurements by calculating the rate of photosynthetic electron transport out of PSII (ETR_PSII_). There are several algorithms that may be applied to derive ETR_PSII_ [17], [26]–[30]. Each algorithm relies on the same principles of light harvesting and photosynthetic electron transport, but arrive at ETR_PSII_ estimates using slightly different, but theoretically equivalent, combinations of photophysiological metrics. As a result, different algorithms confer different field-sampling advantages and challenges (see [31]). The so-called ‘amplitude’ based approach (abbreviated ETR_a_; sometimes also referred to as the sigma-algorithm), calculates ETR_PSII_ as the product of photosynthetically available radiation (PAR), the functional absorption cross section of PSII in the dark-acclimated state (σ_PSII_), and the photochemical efficiency of PSII normalized by the dark-acclimated maximum photochemical efficiency of PSII [27]. This approach reduces uncertainty in ETR_PSII_ estimates by using σ_PSII_ measurements made in the dark-acclimated state, which are subject to less noise than σ_PSII_′ measurements made in the light [31]. As a result, ETR_a_ is a favorable approach in low biomass waters where low signal-noise measurements represent a considerable challenge.

To achieve dark acclimation for ETR_a_ measurements, phytoplankton samples are kept in darkness or very low light to relax non-photochemical quenching processes (NPQ) that upregulate heat dissipation of absorbed light energy, thereby reducing photochemistry and fluorescence yields [32]. Optimal dark acclimation times vary between phytoplankton species, and depend to a large extent on the environmental history of samples [33], making it challenging to design widely applicable field protocols for high resolution data acquisition. In practice, applied dark-acclimation periods range from 5-30 minutes (e.g. [34], [35]). Many FRRf field deployments have focused on discrete sample analysis, applying extended dark acclimation periods to ensure samples reach a short-term steady state condition [25], [36]–[38]. Such discrete sample analysis enables standardized measurements and characterization of light-dependent physiological responses, at the cost of significantly reduced spatial and temporal measurement resolution. In contrast, continuous underway flow-through data acquisition yields high-resolution, real-time measurements of phytoplankton photophysiology, but creates uncertainty in the light exposure history of phytoplankton transiting through a ship’s seawater supply lines. As a result, samples analyzed in continuous mode are neither fully representative of in-situ photophysiology or fully dark-acclimated states, and thus incompatible with the ETR_a_ algorithm that requires both dark- and light-acclimated measurements.

Recently, a new fluorescence approach has been developed to derive ETR_PSII_ based on the turnover rate of the primary electron acceptor molecule (Q_a_) within the photosynthetic electron transport chain [30]. In this ‘kinetic’ approach, Q_a_ turnover rates are derived from analysis of fluorescence relaxation time-constants, resulting in the term ETR_k_. The derivation of ETR_k_ does not depend on dark-acclimated measurements, and can thus significantly increase the frequency of ship-board ETR_PSII_ measurements. The kinetic fluorescence approach for ETR_k_ was originally developed and implemented in mini-Fluorescence Induction-Relaxation (mini-FIRe) instruments [30], which use similar data acquisition protocols, but a different numerical approach for fluorescence relaxation analysis than FRRf. Gorbunov et al. [30] observed greater coherence between growth rates of laboratory cultures and FIRe-derived ETR_k_ compared to alternative ETR_PSII_ estimates, suggesting a strong potential for ETR_k_ to quantify in situ primary productivity. To our knowledge, there have been very few direct comparisons of FRRf-derived ETR_k_ and ETR_a_ estimates for natural phytoplankton assemblages.

Here, we present results from ship-board FRRf analyses quantifying phytoplankton photophysiology and ETR_PSII_ along a ship-track through Lancaster Sound and Barrow Strait in the Canadian Arctic Ocean. We employed a hybrid approach to data collection, combining semi-continuous flow-through measurements with light response curves on static samples. Continuous sampling enabled us to obtain high-resolution measurements and examine the effects of light history and nutrient status on the physiology of Arctic phytoplankton assemblages. Data from rapid light-response curves allowed direct comparison of FRRf-derived ETR_k_ and ETR_a_ estimates. Our results demonstrate residual light-dependent NPQ effects on dark (low light) sample measurements, and illustrate a decoupling of ETR_k_ and ETR_a_ under conditions of phytoplankton photophysiological stress. We relate the spatial patterns in our observations to regional and fine-scale patterns in hydrography and nutrient supply in the eastern Canadian Arctic Archipelago, and discuss the potential effects of different data analysis approaches on ETR_PSII_-based productivity estimates. Results from our work will inform future ship-based deployments of FRRf and related techniques to understand spatial patterns in phytoplankton productivity.

## Materials and Methods

### Underway sampling

Arctic Ocean samples and hydrographic data were collected aboard the *CCGS Amundsen* from August 10 – 15, 2019, within the eastern region of Lancaster Sound and Barrow Strait, in the waters surrounding Devon and Cornwallis Islands. All FRRf measurements were obtained with a Soliense LIFT (Light Induced Fluorescence Transient)-FRR fluorometer (LIFT-FRRf; Soliense Inc., USA). Water samples were delivered to the LIFT-FRRf sampling cuvette using the ship’s seawater line as a primary supply (March MFG Inc, BC-4C-MD pump, nominal flow rate ∼20 L min^-1^), combined with two secondary peristaltic pumps. The first pump (Masterflex L/S, model 7518-10), was used to create a continuous sampling loop (∼200 mL min^-1^) that was connected via t-fitting to a custom-built peristaltic pump actuated by the LIFT-FRRf software. Use of the FRRf-actuated pump enabled precise synchronization of the sample handling and fluorescence measurements, allowing us to employ a semi-continuous sampling strategy of alternating fluorescent transient and light response curve measurements (details below).

In parallel with FRRf measurements, *in-situ* Chl*a* fluorescence, surface water salinity and temperature were measured using a flow-through thermosalinograph system (Seabird, SBE 38), equipped with a Wetlabs fluorometer (WETStar). The underway Chl*a* fluorescence signal was calibrated against discrete samples collected from surface Niskin bottles to approximate the along-track Chl*a* biomass (mg m^-3^). Surface PAR measurements were obtained from an QCR-22000 biospherical probe mounted above the ship’s super-structure. Biological oxygen saturation, ΔO_2_/Ar, was measured as a metric of net community production using a membrane inlet mass spectrometer (MIMS) following the approaches outlined by Tortell et al. [39], [40]. These gas measurements were made continuously on seawater obtained from the same underway lines that supplied the FRRf system. Briefly, seawater was circulated at a constant flow rate past the mass spectrometer’s inlet cuvette consisting of a 0.18 mm thick silicone membrane. Measurements of the mass-to-charge ratios at 32 (O_2_) and 40 (Ar) atomic mass units were obtained at approximately 20 s. intervals. Air standards, consisting of filtered seawater (<0.2 µm) incubated at ambient sea surface temperature and gently bubbled using an aquarium air pump, were run periodically by automatically switching the inflow water source every 45 – 90 minutes. For both seawater and air standard measurements, the inflowing water passed through a 6-m heat exchange coil immersed in a constant-temperature 4°C water bath before passing the MIMS inlet. The seawater and air standard O_2_/Ar ratios ([O_2_/Ar]_sw_ and [O_2_/Ar]_std_, respectively) were used to derive underway ΔO_2_/Ar by linearly interpolating between air standard measurements.

In addition to continuous surface water sampling, water column hydrographic profiles were examined using CTD casts to a depth of 200m at 16 stations (Fig 1). Mixed layer depths were derived from a density difference criterion of 0.125 kg m^-3^ from surface water values.

**Figure 1.**
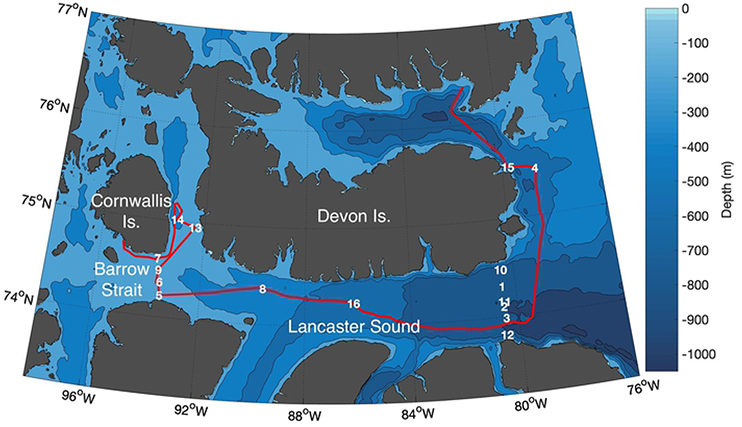
Study Region Map. CTD stations sampled in August 2019 are numbered and overlain on bathymetric contours. We use the 200m bathymetric contour line around 92°W to separate Lancaster Sound and Barrow Strait. The ship track is shown in red.

The CTD and underway surface salinity, temperature, Chl*a*, and PAR data were provided by the Amundsen Science group of Université Laval (Amundsen Science Data Collection, 2020a,b,c).

### LIFT-FRRf Sampling protocols & parameter retrieval

Chlorophyll *a* fluorescence transients were obtained using a Single Turnover (ST) flash protocol and fit to the biophysical model of Kolber et al. [23] to derive photosynthetic parameters under dark acclimation and under actinic light, where the latter is denoted with the’ notation. A summary of parameters measured and definitions is given in Table 1. Specifically, we derived estimates of the functional absorption cross section of PSII (σ_PSII_, σ_PSII_′), the minimum and maximum fluorescence when the reaction center pool is fully open and closed (*F_o_*, *F*’ and *F_m_*, *F_m_*’, respectively), and the variable fluorescence (*F_v_* = [*F_m_* - *F_o_*]; *F_q_*’ = [*F_m_*’ - *F*’]). Single Turnover flash excitation flashlets were delivered simultaneously by 445, 470, 505, 535, and 590 nm LED lamps. Each curve fit and acquisition was based on fluorescence yields averaged from five sequential ST flash sequences delivered every 1s. Curve fitting and fluorescence averaging was completed within Soliense LIFT software.

**Table 1.**
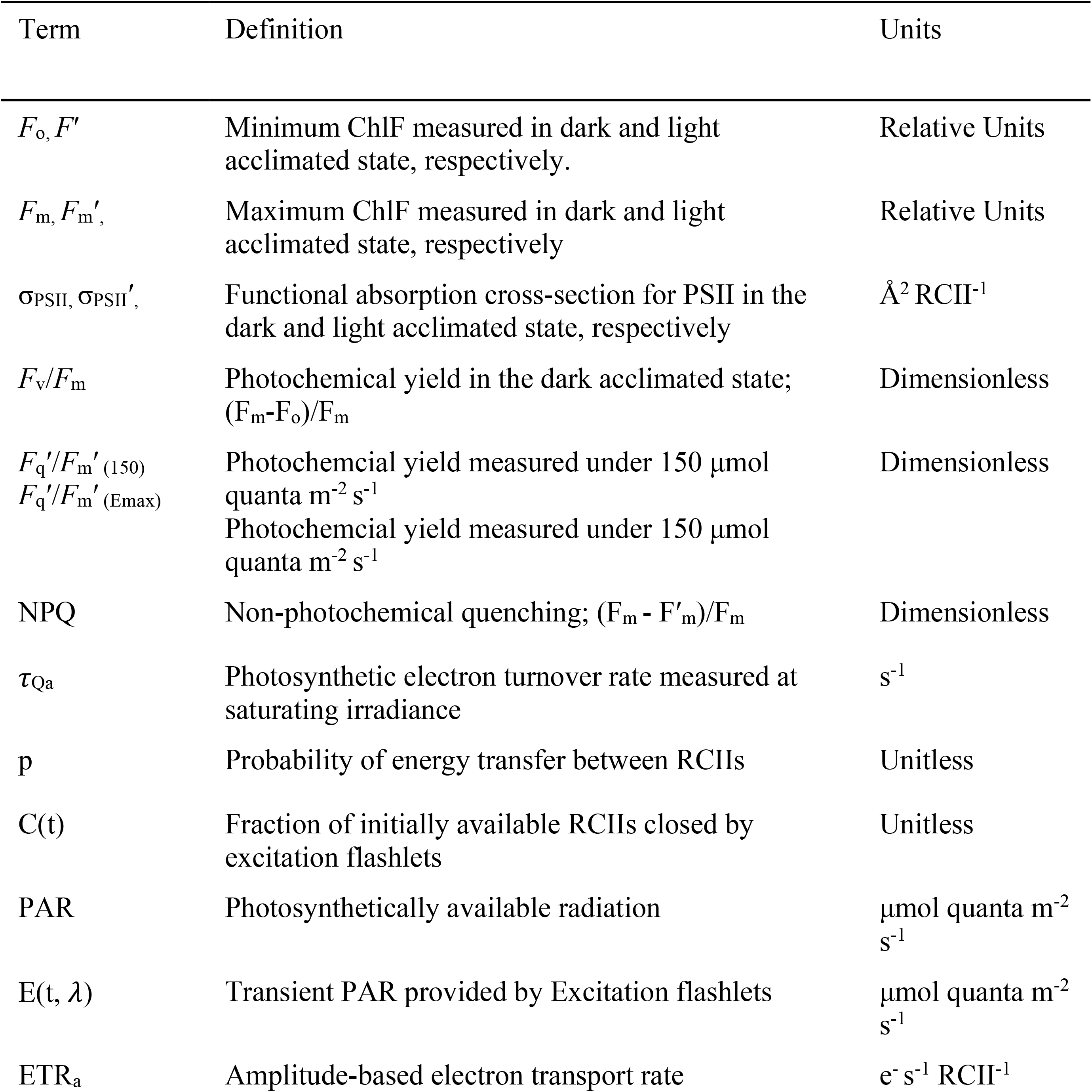
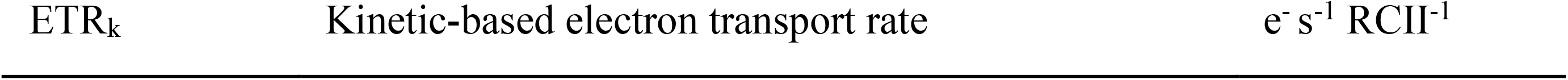
Commonly referred to photophysiological terms and abbreviations.

Each seawater sample was initially held in the cuvette under low light for five minutes to allow NPQ relaxation. During this low-light acclimation period, five LEDs at peak wavelength excitation of 445, 470, 505, 535, and 590 nm each supplied 1 μE m^-2^ s^-1^ of actinic light, providing a total irradiance of 5 μE m^-2^ s^-1^. Samples were acclimated to low light rather than total darkness to avoid RCII closure and fluorescence quenching associated with back flow of electrons from the PQ pool to Q_a_ (e.g. [41]). After this short acclimation period, 40 acquisitions (each consisting of 5 averaged fluorescence transients) were collected. Irradiance was then increased to 150 μE m^-2^ s^-1^ (30 μE m^-2^ s^-1^ per LED) and samples were held for an additional five minutes to acclimate to the higher irradiance level before collecting another 40 acquisitions. The LIFT-FRRf cuvette was then flushed for 60 s with seawater from the ship seawater supply, displacing at least five full cuvette volumes before the pump was turned off to isolate the next sample for measurements.

Semi-continuous sampling was interrupted every 90 mins to perform light response curves on static samples. For each light response curve, the actinic irradiance supplied by each LED lamp increased incrementally from 0 to 350 μE m^-2^ s^-1^ for all 5 LEDs to create light steps of 0, 15, 35, 75, 110, 150, 200, 300, 400, 550, 850, 1250, and 1750 μE m^-2^ s^-1^ total actinic light. A total of 25 acquisitions (of 5 averaged fluorescence transients each), were obtained at each light step. Fluorescence amplitudes generally stabilized by the third round of data acquisition, and we thus excluded the first two flash sequences at each light level from data analysis. Light response curves were used to calculate ETR_PSII_ as a function of increasing light intensity (see below for calculation details). To produce final photosynthesis-irradiance curves, ETR_PSII_ was plotted against irradiance and fit to the hyperbolic function described by Webb et al. [42], using a least squares non-linear regression to derive the maximum light-saturated photosynthetic rate (ETR_max_), the saturating light intensity (E_k_) and the light utilization efficiency (α).

Daily blank corrections were performed by analyzing seawater gently passed through a 0.2 μm syringe filter (e.g. [25]). Soliense software was programmed to subtract the resulting fluorescence intensity from all measurements. Prior to the field deployment, the LIFT-FRRf lamps used to produce actinic light and the probing flashes were calibrated using a Walz spherical submersible micro quantum sensor (Walz, US-SQS-L).

### Electron Transport Rates (ETRPSII)

The amplitude and kinetics-based ETR_PSII_ algorithms (ETR_a_ and ETR_k_, both e^-^ s^-1^ RCII^-1^) were applied to fluorescence data collected during light response curves to determine ETR_PSII_ at increasing actinic light intensities. ETR_a_ was applied to all 85 light response curves collected along the ship-track to produce photosynthesis-irradiance curves following Webb et al. [42]. At each light level, ETR_a_ (e^-^ RCII^-1^ s^-1^) was determined as the quantity of incident light absorbed by PSII directed towards photochemistry [27], [28];

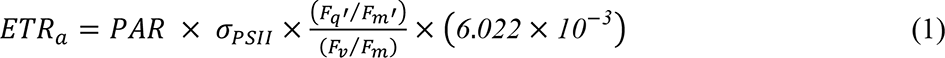

Here, PAR (μmol quanta m^-2^ s^-1^) is the total actinic light provided by the FRRf LEDs, σ_PSII_ (Å^2^ quanta^-1^) is the PSII functional absorption cross section measured in dark acclimated samples, and *F_q_ʹ*/*F_m_*ʹ divided by *F_v_*/*F_m_* (dimensionless) is the PSII photochemical efficiency measured under actinic light normalized by the dark-measured maximum photochemical efficiency. The constant 6.022*10^-3^ converts σ_PSII_ units from Å^2^ RCII^-1^ to m^2^ RCII^-1^ and PAR from *μ*mol quanta to quanta.

For ETRk calculations, fluorescence data from a subsample of 25 light response-curves with optimal signal-to-noise ratios were re-analyzed using a 3-component multi-exponential model to describe Q_a_ reoxidation kinetics. This numerical procedure was applied as a fitting option in the FRRf Soliense Software.

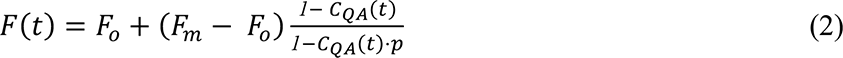

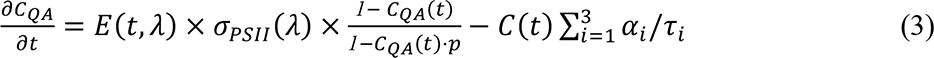

Here, C(t) (dimensionless) is the fraction of available PSII reaction centers closed by excitation flashlets, with C(t) equal to 1 when the photochemistry pathway is fully closed. The term p (dimensionless) is the connectivity factor, which describes the likelihood of energy transfer between RCIIs. *α*_5_ and *T*_5_ refer to the amplitude and time constant of the i^th^ component of Q_a_ reoxidation, respectively. The value 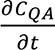 is determined by the balance between primary photochemistry induced by excitation flashlets (E; μmol quanta m^-2^ s^-1^) and electron transfer from Q_a_ to secondary electron acceptor, Q_b_, mediated by three kinetic components. Fluorescence transients were fit to retrieve photophysiological parameters by integrating Eq. 3 over the length of the ST flash sequence and then iteratively fitting Eq. 2. to the fluorescence data. Importantly, we note Eq. 3 differs slightly from that applied by FIRe-based data analysis, in which there is an added term to describe RCII closure induced by actinic irradiances (See*‘Computational Considerations’* in the Discussion).

The rate of Q_a_ reoxidation was calculated by averaging the time constants of the two primary components of electron transfer from Q_a_ to secondary electron acceptor Q_b_ following the approach of Gorbunov and Falkowski [30] as,

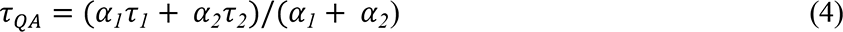

ETR_k_ was then derived at each light step as

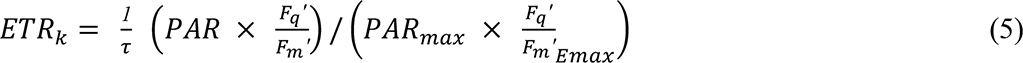

Here, PAR_max_ is a super-saturating light level chosen as a value three–fold higher than the light saturation parameter, E_k_, derived from photosynthesis irradiance curves. 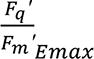 is the PSII photochemical efficiency measured under PAR_max_. The photosynthetic turnover rate, 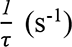, is taken as 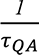 at saturating irradiance where primary photochemistry is at a maximum [30].

### Non-Photochemical Quenching (NPQ)

We quantified NPQ as a measure of the relative increase in heat dissipation of absorbed energy by PSII between samples exposed to low light and 150 *μmol quanta* m^2^ s^-1^, following Bilger and Bjorkman [43] as:

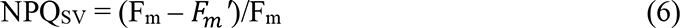

This derivation assumes full relaxation of all NPQ processes in the dark-acclimated state. However, as discussed in Results section, this assumption did not hold in our samples. For this reason, we used an additional measure of long-term NPQ processes, based on the ratio of photochemical efficiency measured under low and high light (i.e. (*F*_q_*ʹ* /*F_m_ʹ*)/(F_v_/F_m_)).

### Statistical analysis

Pair-wise statistical relationships between measured variables were determined using Spearman Rank correlation tests. Samples from two regions of our transect (Lancaster Sound and Barrow Strait) were compared using Kruskal-Wallis test (Fig 8). Lilliefors test rejected the null hypothesis that underway data were normally distributed, so we report median rather than mean values of all photophysiological variables. Deviation from the median was determined as the median absolute deviation. Continuously measured, in-situ parameters such as temperature and salinity were not corrected for autocorrelation. All analyses were completed using Matlab (R2020a).

## Results

### Hydrographic properties

Sea surface temperature (SST) and salinity varied significantly across our study region, reflecting the influence of different water masses and freshwater inputs. Sea surface temperature ranged from 0.6 to 15.5 °C, and salinity ranged from 22.8 to 31.6 PSU. Salinity and SST strongly covaried, and exhibited a number of sharp transitions across prominent hydrographic fronts associated with freshwater input (Fig 2; Table 2). Surface layer (∼7m) phytoplankton biomass, approximated by *in-situ* Chl*a* fluorescence measurements (non-FRRf), was low throughout the entire transect, with a mean Chl*a* fluorescence equivalent to 0.15 ± 0.04 mg m^-3^ (*n*= 7200). These in situ Chl*a* measurements showed significant diel periodicity, likely reflecting daytime quenching effects. Both SST and salinity exhibited weak (though statistically-significant) correlations with Chl*a* (Table 2). Mixed layer depths recorded at the 16 profiling stations were shallow, with a mean of 9.2 ± 4.5 m. Mixed layer nitrate and nitrite concentrations were near zero at each of the depth profiling stations (Table 3, mean 0.03 ± 0.049 *μ*M, *n* = 16), indicative of post-bloom, nutrient-limited summer conditions. The mixed layer Chl*a* concentration averaged across all 16 profiling stations was 0.40 ± 0.18 mg m^-3^ (*n* = 16). There was no statistically significant relationship between station Chl*a* and mixed layer nitrate and nitrite concentrations (*r* = 0.09, *p* = 0.75, *n* = 16). Biological oxygen saturation, derived from ΔO_2_/Ar measurements, was greater than zero across the entire cruise track, implying net autotrophic conditions. Small-scale features in ΔO_2_/Ar distributions were observed across hydrographic frontal regions with only weak correlation to salinity or temperature measured along the cruise track (Table 2).

**Fig 2.**
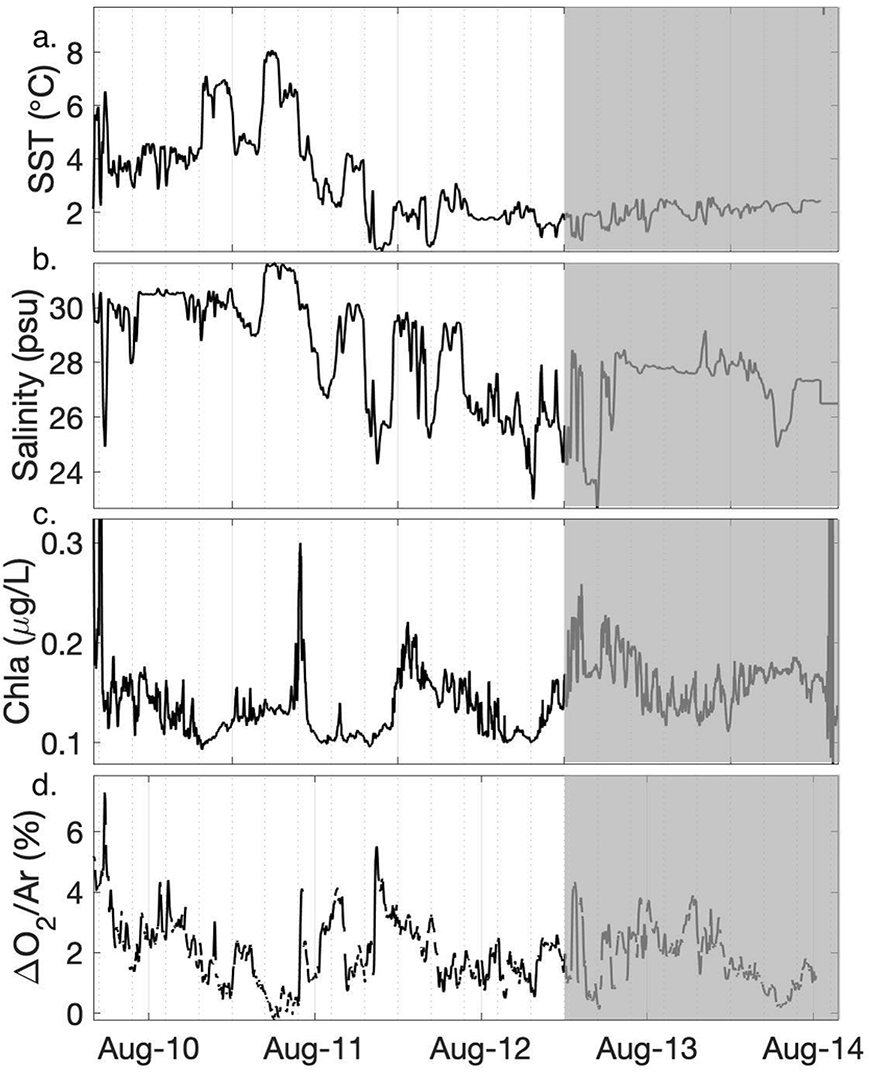
Summary of oceanographic conditions. Hydrographic conditions, phytoplankton biomass and biological oxygen saturation measured in surface waters (∼ 7 m depth) along the cruise track. Shaded area on Aug. 13-14 marks measurements made in Barrow Strait.

**Table 2.**
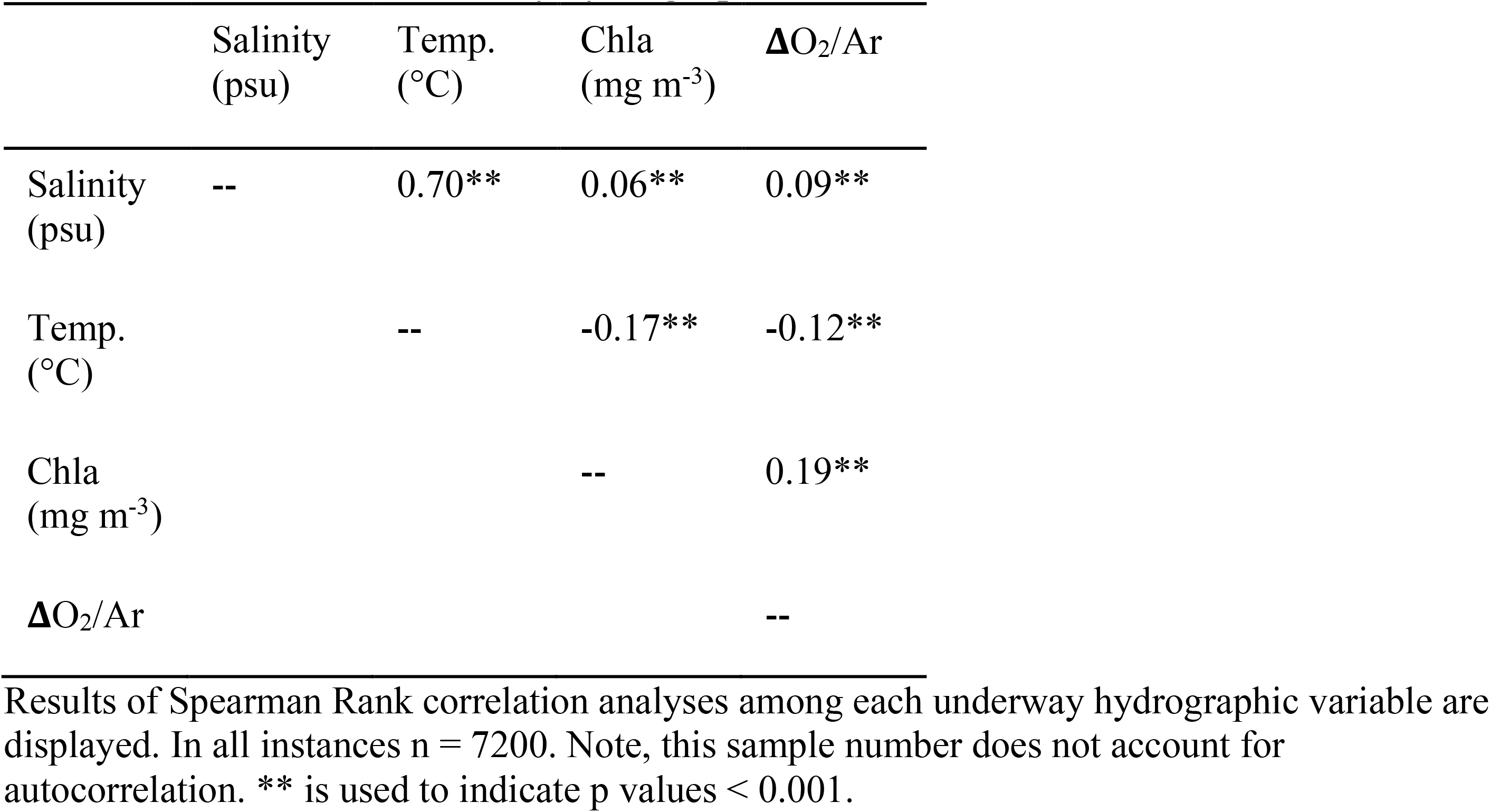
Correlation of underway hydrographic variables.

| | Salinity<br>(psu) | Temp.<br>(°C) | Chla<br>(mg m<sup>-3</sup>) | $\Delta O_2/Ar$ |
| --- | --- | --- | --- | --- |
| Salinity<br>(psu) | -- | 0.70** | 0.06** | 0.09** |
| Temp.<br>(°C) |  | -- | -0.17** | -0.12** |
| Chla<br>(mg m<sup>-3</sup>) |  |  | -- | 0.19** |
| $\Delta O_2/Ar$ | | | | -- |

Results of Spearman Rank correlation analyses among each underway hydrographic variable are displayed. In all instances $n = 7200$ . Note, this sample number does not account for autocorrelation. \*\* is used to indicate $p$ values $< 0.001$ .

**Table 3.** CTD profiling stations.

| Location | CTD<br>Station | Latitude<br>(°) | Longitude<br>(°) | MLD<br>(m) | [NO<sub>3</sub><sup>-</sup> +<br>NO<sub>2</sub><sup>-</sup>] (μM) | Chl *a*<br>(mg/m<sup>-3</sup>) |
| --- | --- | --- | --- | --- | --- | --- |
| LS | 1 | 74.316 | -80.517 | 6 | 0.002 | 0.43 |
| LS | 2 | 74.107 | -80.511 | 9 | 0.007 | 0.21 |
| LS | 3 | 73.987 | -80.468 | 9 | 0 | 0.25 |
| LS | 4 | 75.482 | -78.643 | 8 | 0 | 0.12 |
| BS | 5 | 74.222 | -93.517 | 10 | 0.007 | 0.20 |
| BS | 6 | 74.361 | -93.556 | 6 | 0.003 | 0.39 |
| BS | 7 | 74.599 | -93.717 | 10 | 0.013 | 0.41 |
| LS | 8 | 74.371 | -89.621 | 9 | 0.087 | 0.50 |
| BS | 9 | 74.483 | -93.640 | 9 | 0.063 | 0.77 |
| LS | 10 | 74.486 | -80.508 | 8 | 0.014 | 0.45 |
| LS | 11 | 74.159 | -80.470 | 6 | 0.189 | 0.24 |
| LS | 12 | 73.816 | -80.508 | 7 | 0 | 0.58 |
| BS | 13 | 74.951 | -92.388 | 24 | 0.018 | 0.58 |
| BS | 14 | 75.014 | -93.115 | 14 | 0.048 | 0.28 |
| LS | 15 | 75.523 | -79.748 | 12 | 0.047 | 0.54 |
| LS | 16 | 74.234 | -86.160 | 6 | 0 | 0.21 |

Sampling station locations are indicated as LS for Lancaster Sound or BS for Barrow Strait. Station mixed layer depth (MLD), mean mixed layer nitrate and nitrite concentration, and Chl *a* concentration within the mixed layer are shown.

### Photophysiological measurements

#### Continuous flow-through measurements

The photophysiological parameters *F_v_*/*F_m_* and σ_PSII_ fluctuated over diel cycles throughout our sampling period (Fig 3a-b), showing daily maxima during the night, and decreasing during daylight hours. Both of these variables exhibited a significant negative correlation with actinic surface PAR intensity averaged over the 5 min window prior to sample measurements (*r* = −0.50 and *r* = −0.72, respectively, *p* < 0.001 and *n* = 481 for both; Fig 4a-b). These observations are consistent with previously noted *in-situ* photophysiological diurnal patterns of daytime fluorescence quenching [44], [45]. We note, however, that these *F_v_*/*F_m_* and σ_PSII_ measurements were collected after 5 minutes of low light exposure, indicating the persistence of longer-lived light-dependent quenching effects after this acclimation period. We thus conclude that dark-acclimation (i.e. NPQ relaxation) was not achieved in our measurement protocol, and *F_v_*/*F_m_* and σ_PSII_ are thus representative of an intermediate state between *in-situ* and dark-acclimated values.

**Fig 3.**
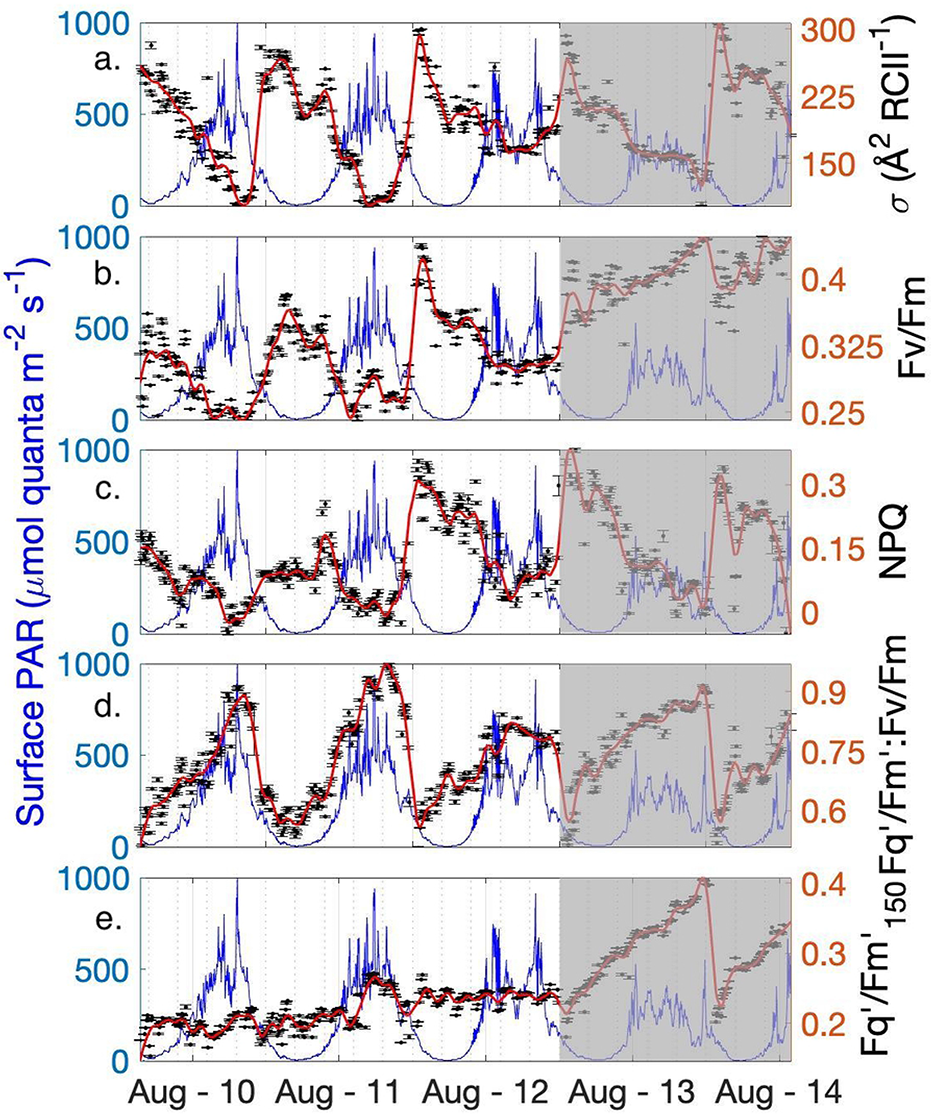
Photophysiological conditions summary. Semi-continuous FRRf measurements of photophysiology (black dots) are superimposed over the in-situ surface PAR (blue line). A loess smoothing function was applied to photo-physiological measurements (red line). Shading denotes the Barrow Strait portion of the transect.

**Fig 4.**
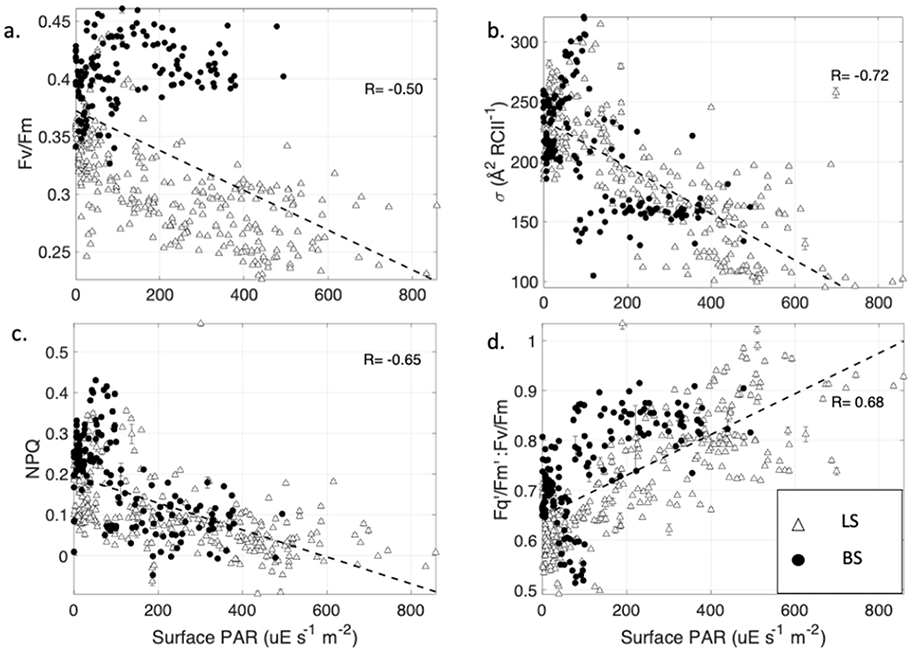
Recent light history effects on photophysiology measured under low light. FRRf-derived photophysiological parameters measured under low light are plotted against in-situ surface PAR at the time of sample acquisition. Panels (a) and (b) show Fv/Fm and σ_PSII_ measurements made after 5 min of low light exposure. Panels (c) and (d) show NPQ and residual quenching measured after low light treatment. Data collected in Barrow Strait (BS) are shown in filled points, while data collected in Lancaster Sound (LS) are represented with unfilled circles. Error bars show standard error, but are often concealed by size of data symbols. R values indicate Spearman Rank correlation coefficients.

Despite signs of strong light-dependent quenching, NPQ values, calculated as NPQ_SV_ = (F_m_ – F_m_ʹ)/F_m_, were negatively correlated to surface PAR (Fig 4c). This surprising result can be explained by the derivation used for here for NPQ, which measures the fractional change in NPQ between light and dark (low light) measurements, rather than total NPQ. As a result, this approach does not account for any residual quenching present in samples after five minutes of low light acclimation. To address this limitation, we used the ratio (F ʹ_q_/F ʹ_m_)/(F_v_/F_m_) to estimate the extent of residual quenching present in samples after five minutes of NPQ relaxation. As expected, this derived variable was well correlated to surface PAR (Fig 4d, *r* = 0.68, *p* < 0.001, *n* = 481), reflecting the effects of residual quenching and, potentially, short-term photoinhibition.

To further examine low light-acclimated *F_v_*/*F_m_* and σ_PSII_ values, we isolated night-time *F_v_*/*F_m_* and σ_PSII_ measurements collected under relatively low ambient surface PAR. Due to the long summer daylight hours in the Arctic, only 1% of data points represents true night when surface PAR = 0. We thus chose light levels ≤100 μmol quanta m^-2^ s^-1^ to represent night-time conditions. By comparison, midday surface PAR ranged from 400 – 1000 μE m^-2^ s^-1^. The median night-time *F_v_*/*F_m_* for the entire transect was 0.36 ± 0.03 (*n* = 214), a value similar to the global average of 0.35 ± 0.11 [46], but lower than the 0.55 median value previously recorded for late-summer assemblages in the Canadian Arctic [47]. The lowest *F_v_*/*F_m_* values were recorded at the beginning of the ship track (August 10-12) within Lancaster Sound (0.32 ± 0.03, *n* = 120), while *F_v_*/*F_m_* increased significantly in Barrow Strait (0.40 ± *0*.*02*, *n* = 94), indicating greater photosynthetic potential in this region (Fig 3b). In contrast, night-time σ_PSII_ did not significantly vary between Lancaster Sound (250 ± 17.2) and Barrow Strait (241.1 ± 15.3) (Fig 3a). These absolute σ_PSII_ values are somewhat lower than those reported in previous studies, likely reflecting our use of simultaneous excitation flashlets centered around 445, 470, 505, 535, and 590 nm. Relative to blue light, not all of these wavelengths are efficiently absorbed by phytoplankton, resulting in an apparent decrease σ_PSII_ [48]. As discussed below, σ_PSII_ values are also subject to physiological, taxonomic and environmental effects [49].

As observed for *F_v_*/*F_m_*, the photosynthetic efficiency measured under 150 *μmol quanta s,^1^m,^2^* actinic light (*F_q_ʹ*/*F_m_*ʹ_(150)_) displayed strong regional differences, increasing from 0.21 ± 0.01 (*n* = 120) in Lancaster Sound to 0.30 ± 0.02 (*n* = 94) in Barrow Strait. However, unlike the low-light measurements, *F_q_ʹ*/*F_m_*ʹ_(150)_ did not exhibit a diel signature and was remarkably consistent across samples measured within Lancaster Sound (Figs 3e and 5). Moreover, there was no relationship between *F_q_ʹ*/*F_m_*ʹ _(150)_ values and surface PAR intensity experienced prior to sampling (*r* = 0.01, *p* = 0.76, *n*= 481). This result suggests that irradiance-specific PSII photochemical yields were independent of prior light-history experienced by phytoplankton. This, in turn, indicates that even short (5 min) exposure to actinic light is sufficient for phytoplankton to reach a photosynthetic steady-state.

**Fig 5.**
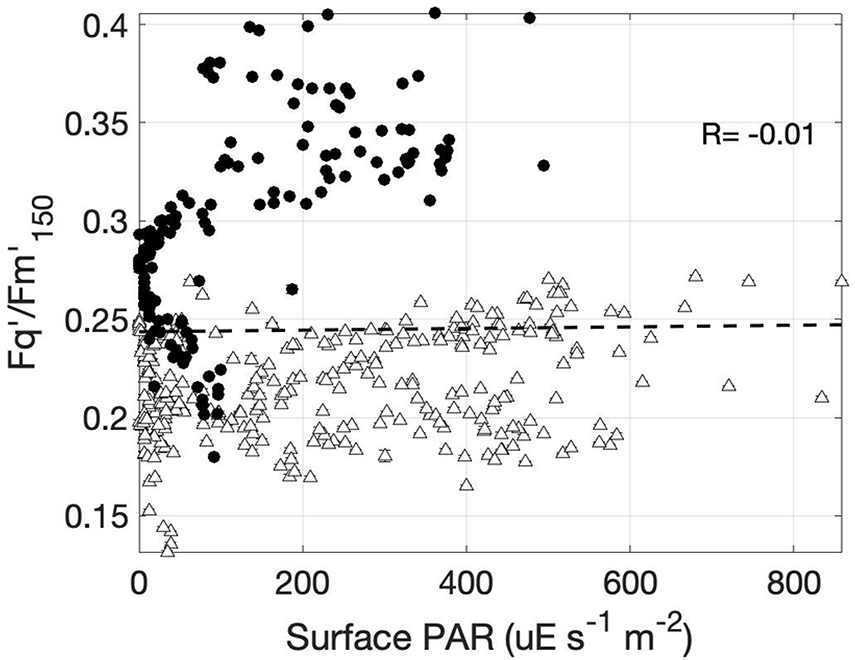
Recent light history effect on photochemical efficiency measured under constant actinic light. The PSII photochemical efficiency measured under 150 μE m^-2^ s^-1^ in relation to natural surface irradiance at the time of sample acquisition. R value indicates Spearman Rank correlation coefficient. Data collected in Barrow Strait (BS) is shown in filled points, while data collected in Lancaster Sound (LS) is represented with unfilled circles.

### Photosynthesis-Irradiance curves and ETRPSII comparisons

We used light response curves to compare FRRf-based ETR_a_ and ETR_k_ estimates. In this approach, ETR_a_ (Eq. 1) was plotted against actinic irradiance (Fig 6) to derive the maximum rate of charge separation at RCII (ETR_max_), the light-dependent increase in the charge separation rates (*α*), and the saturating light intensity (E_k_). Fit parameters from these curves varied considerably, with mean values of ETR_max_, *α*, and E_k_ of 460 ± 345 e^-^ s^-1^ RCII^-1^, 1.37 ± 0.57 e^-^ RCII^-1^ quanta^-1^ m^-2^, and 358.0 ± 199.0 μmol quanta m^-2^ s^-1^, respectively (*n* = 85).

**Fig 6.**
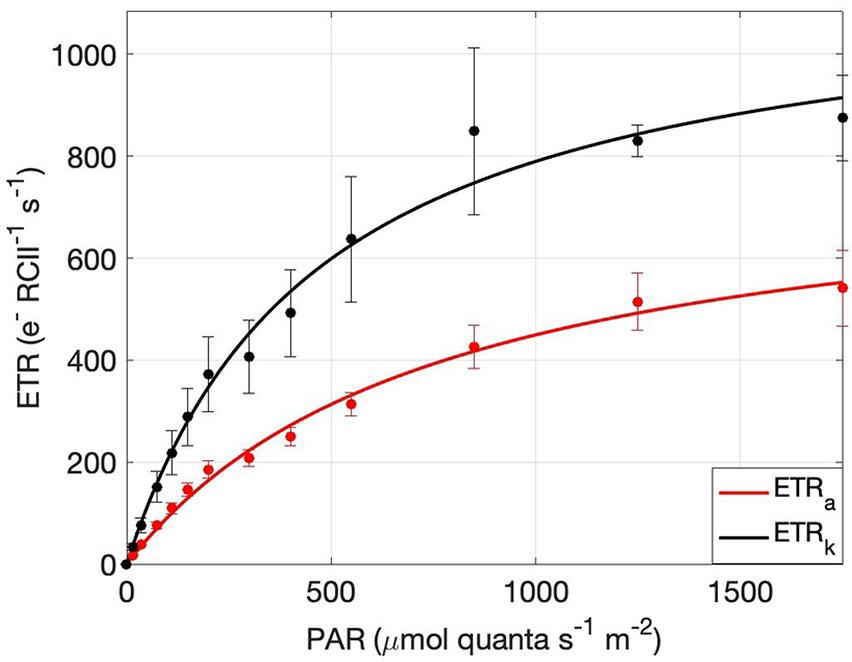
Photosynthesis-Irradiance curves derived by ETR_k_ and ETR_a_. Consolidated mean ETR_k_ (black) and ETR_a_ (red) estimates at each light step of the 25 reprocessed P-I curves. Error bars represent the standard error of all individual measurements from all curves at each light step. Curves were produced using the photosynthesis-irradiance function described by Webb et al. (1974).

A subsample of 25 light response curves was re-analyzed using the ‘kinetic’ approach (ETR_k_; Eq. 5), and compared with ETR_a_ values. This comparison revealed a strong correlation between the ETR values (*r* = 0.81, *p* < 0.001; Fig 7). However, the kinetics-based algorithm produced consistently higher results than ETR_a_, with values 1.96 ± 1.2 times greater, on average, than ETR_a_ (Fig 6).

**Fig 7.**
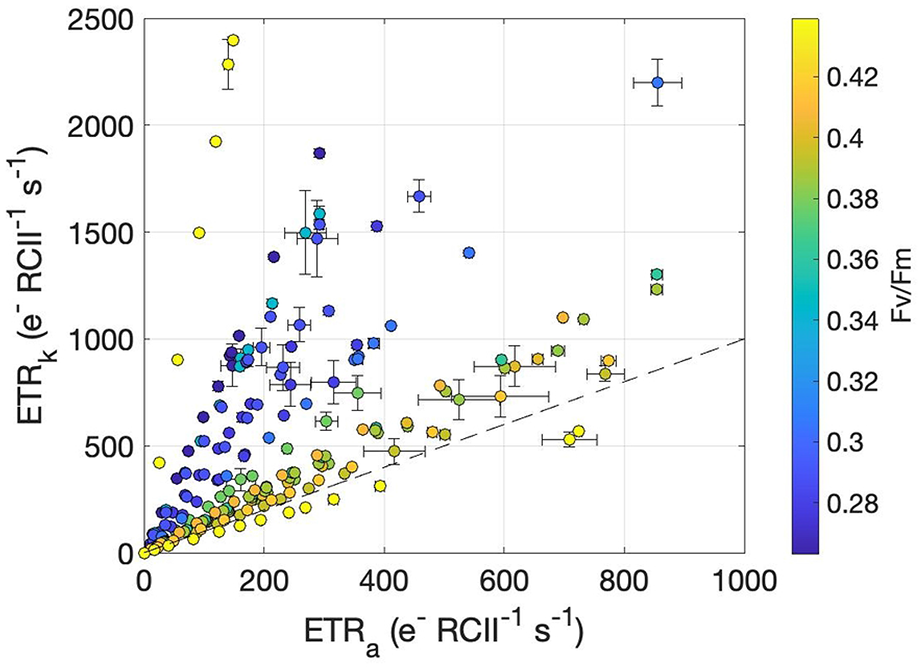
ETR_k_ plotted against ETR_a_ derived from Photosynthesis-Irradiance curves and colored by Fv/Fm. Each point represents the mean ETR value at a given light intensity within a Photosynthesis-Irradiance curve, and error bars are the standard error. The dashed line indicates a 1:1 relationship.

Along our cruise track, we observed a distinct spatial pattern in the ETR_k_:ETR_a_ ratio. The highest values (3.26 ± 0.95, *n* = 14) occurred during the early part of our survey (August 10-13), with significantly lower ETR_k_:ETR_a_ (1.42 ± 0.16, n = 11) observed near the end of our cruise, particularly in Barrow Strait where F_v_/F_m_ was maximum (Fig 8). More generally, ETR_k_:ETR_a_ displayed a strong negative correlation with *F_v_*/*F_m_* values (*r* = −0.60, *p* < 0.01, *n* = 25). As discussed below, this result suggests ETR_k_:ETR_a_ decoupling is strongly driven by photophysiological and environmental variability.

**Fig 8.**
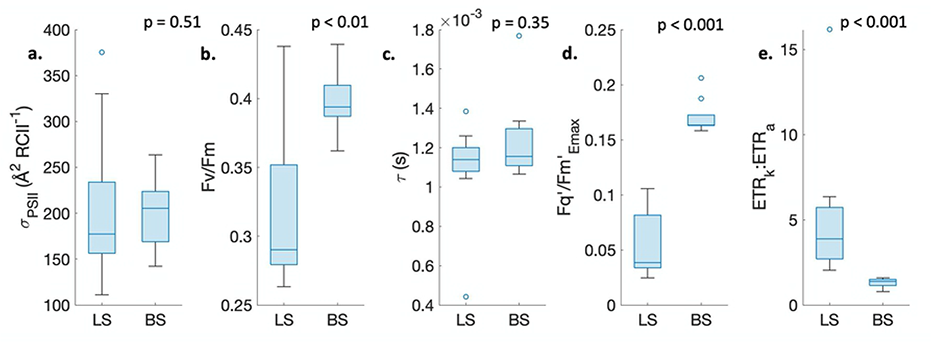
Parameters contributing to the ETR_k_:ETR_a_ ratio measured in Lancaster Sound (LS) versus Barrow Strait (BS). Horizontal lines within each boxplot represent the median. The upper and lower edge of each box demarks upper and lower quartiles, respectively, while whiskers extend over the entire data range, excluding outliers. Outliers, determined as data points falling over 1.5x the interquartile distance away from box edges, appear as unfilled circles. P values in each subplot are results from 2-group Kruskal-Wallis tests. All data shown here was collected during the 25 reprocessed P-I curves.

## Discussion

The primary focus of our work was to quantify phytoplankton photophysiology and productivity along our ship-track using active Chl*a* fluorescence methods. With this in mind, we applied a sampling and analysis strategy to support both amplitude-based and kinetic analysis of FRRf data to derive ETR estimates. In the following, we first discuss the spatial patterns in FRRf-based measurements in relation to nutrient concentrations and light histories of phytoplankton assemblages encountered along our sampling transect through Lancaster Sound. We then examine potential factors leading to the uncoupling of ETR_PSII_ estimates, including environmental and physiological variability, and potential influences of different data analysis methods. We conclude by discussing the implications of our results for future ship-board FRRf deployments.

### Spatial distribution of photophysiology

Across our survey region, we observed notable spatial patterns in FRRf data, with low *F_v_*/*F_m_* and *F_q_ʹ*/*F_m_*ʹ_(150)_ values in Lancaster Sound, and significantly higher values in Barrow Strait. Previous studies of late-summer Arctic photophysiology have reported low background *F_v_*/*F_m_* values, as observed over most of our sampling transect. At the same time, these prior studies have also demonstrated increased *F_v_*/*F_m_* in response to nitrate (but not phosphate) enrichments [5], [50]. This result, coupled with the nutrient depletion observed at profiling stations along our cruise track (Table 2), suggest that the low *F_v_*/*F_m_* values we observed likely reflect nitrogen deficiency. Iron (Fe)-limitation has also been shown to exert a strong negative effect on *F_v_/F_m_*, coincident with increases in σ_PSII_ values. These Fe-dependent effects result from a combination of physiological responses [36], [51], [52] and taxonomic shifts towards smaller cells [49]. In contrast, the low *F_v_*/*F_m_* values recorded in the Lancaster Sound were not associated with high σ_PSII_, and correlation analysis of night-time σ_PSII_ and *F_v_*/*F_m_* values revealed a weak positive relationship between σ_PSII_ and *F_v_*/*F_m_* (*r* = 0.17, *p* < 0.05, *n* = 214). Based on these results, and the proximity of our sampling to land-based Fe sources [53], we infer that nitrogen, rather than iron deficiency was the most likely cause of low photo-efficiencies observed from August 10-13 in Lancaster Sound.

Notwithstanding the low background nutrient concentrations measured at profiling stations, chemical analyses from the Canadian Arctic Archipelago Rivers Program and Canadian Arctic GEOTRACES program have revealed elevated nitrite and nitrate concentrations within several rivers that discharge into Barrow Strait (Fig 9; [11], [54]). In this region, we observed low surface water salinity and elevated *F_v_*/*F_m_*, suggesting a link between river input and increased photo-efficiency, which we ascribe to nutrient inputs. Additionally, the greatest mixed layer depths were found at the two CTD profiling stations situated between Cornwallis Island and Devon Island (Table 2), indicative of enhanced mixing associated with strong tidal currents, in agreement with model predictions of elevated mixing in Barrow Strait. These observations suggest that spatial differences observed in FRRf-derived photophysiology may reflect elevated nutrient availability in Barrow Sound, resulting from a combination of river inputs and mixing effects.

**Fig 9.**
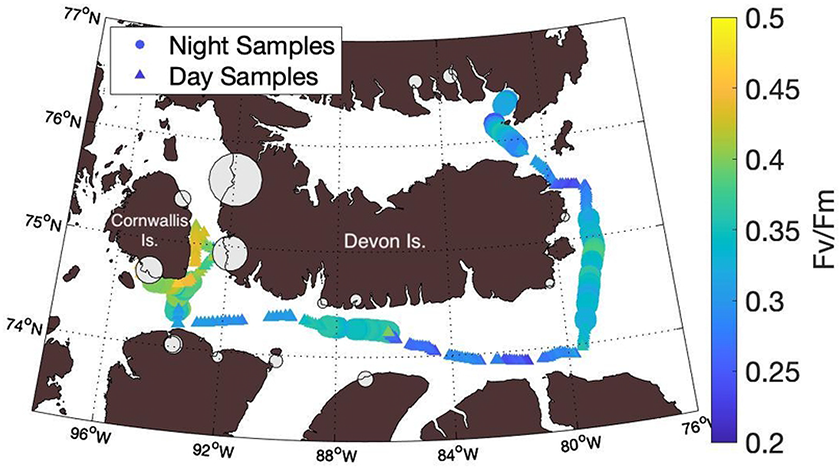
Spatial distribution of riverine nutrient inputs and photochemical efficiency (Fv/Fm) along the ship track. River contributions of nitrate and nitrite are indicated by the size of grey bubbles. The largest inputs of nitrate and nitrite in the region are concentrated in the strait between Cornwallis and Devon Islands, coincident with observations of raised Fv/Fm values. Larger circles are used to denote night-time measurements of Fv/Fm, whereas smaller triangles denote day-time measurements.

Several mechanisms can explain the apparent effects of increased nitrogen availability on phytoplankton photophysiology. First, increased nitrogen availability enables protein synthesis needed to repair inactive reaction centers [55], [56]. Second, localized nutrient loading may indirectly affect photophysiology by stimulating a shift from small to larger phytoplankton species, for instance from nano-flagellates to diatoms [50]. Unfortunately, we lack information on phytoplankton assemblage composition, and thus cannot directly examine any potential taxonomic effects on photophysiology. However, such a shift from small to large cells would be expected to drive a decrease in σ_PSII_, concurrent with increases in *F_v_*/*F_m_* [49], which we did not observe. Previous pigment analyses conducted in the Canadian Arctic Archipelago found that Lancaster Sound and Barrow Strait were both dominated by diatom species, followed by dinoflagellates, in summer [57]. We thus infer that the spatially-divergent *F_v_*/*F_m_* values, coupled with persistently low σ_PSII_, primarily reflect photophysiological nutrient effects in relatively large cells.

### Light-dependent effects and residual NPQ

We observed strong residual NPQ effects after five minutes of low light acclimation (Figs. 3, 4). Notably, the extent of these quenching effects was a predictable function of the short-term light history experienced by in situ phytoplankton assemblages (Fig 3d). Previous studies examining the drivers of NPQ variability [58]–[60] suggest that the magnitude of NPQ effects at a given light level is tied to a number of environmental factors (e.g. temperature and CO_2_ concentrations), phytoplankton taxonomy and physiological status. Given these sources of variability, NPQ relaxation times needed for robust *F_v_*/*F_m_* measurements are expected to differ significantly across ocean regimes. In cold waters, such as those encountered along our ship track, NPQ relaxation is slower [61], and this may have contributed to the longer-lived quenching observed in our low-light samples. As the spatial and temporal resolution of *F_v_*/*F_m_* and ETR_a_ measurements are constrained by such acclimation periods, it is recommended that future field deployments of FRRf conduct experiments using natural assemblages to determine the regional minimum relaxation period necessary to achieve steady-state dark-acclimation. This acclimation step can then be incorporated into underway FRRf protocols, resulting in more robust ETR_a_ estimates, albeit with reduced measurement frequency. Such routine determinations of the minimum NPQ relaxation time requirements have not been commonly carried out for marine phytoplankton [25]. As a result, there is little systematic knowledge of the global variability of NPQ relaxation times. Adopting such pre-study tests (or within protocol) as standard practice would improve current understanding of environmental and taxonomic controls on NPQ relaxation kinetics. Moreover, it may be necessary to adjust the length of the dark-acclimation period to reflect changing conditions over the duration of a cruise. We thus recommend that future work incorporate semi-regular assessments of dark-acclimation times into field-sampling protocols.

## Decoupling of ETRa and ETRk

We observed significantly higher ETR_k_ compared to ETR_a_ across our study region (Fig 6), with the magnitude of ETR_k_ and ETR_a_ decoupling varying strongly in response to phytoplankton photophysiological conditions (Fig 7). The ratio between ETR_a_ and ETR_k_ depends on a number of variables:

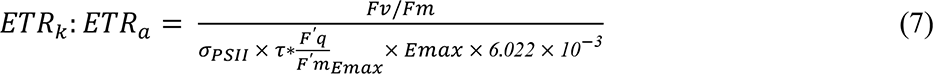

Importantly, all of the terms defining ETR_k_:ETR_a_ are potentially responsive to shifts in nutrient abundances and phytoplankton taxonomic composition. Nutrient enrichment experiments have demonstrated increasing *τ*_Qa_ with nutrient deficiency and elevated actinic irradiance [30], [50]. Moreover, *σ_PSII_* can also change with nutrient availability, but the observed percent change in *σ_PSII_* following short-term nitrate enrichment is small compared with that of *F_v_*/*F_m_* [5], [50]. Baseline fluorescence, may also influence the terms used to define ETR_k_:ETR_a_. This phenomenon represents a non-variable contribution to the Chl fluorescence signal, which is understood to reflect the presence of energetically decoupled light harvesting complexes under nutrient limitation or photoinhibitory stress [31]. High baseline fluorescence decreases the amplitude of fluorescence transients, making *F_v_*/*F_m_* a useful gauge of phytoplankton physiological stress [5], [62], [63]. Analysis of our data revealed a strong correlation between *F_v_*/*F_m_* and (F’q/F’m)_Emax_, (*r =* 0.93, *p* < 0.001, *n*= 25), suggesting (F’q/F’m)_Emax_ is similarly affected by baseline fluorescence and reflective of physiological status. Overall, we thus conclude that differential environmental and taxonomic sensitivities of the variables used to derive ETR_a_ and ETR_k_ can lead to discrepancies between these two productivity metrics.

Going forward, further investigation of ETR_k_ and ETR_a_ divergence is critical to inform our understanding of electron requirements for carbon assimilation and biomass production, particularly under nutrient limiting conditions [28]. For instance, Schuback et al. [36] found that iron-limited phytoplankton assemblages exhibited elevated ETR_a_ and greater decoupling between ETR_a_ and C-assimilation rates as compared to iron-enriched assemblages. This result was attributed to the higher σ_PSII_ values in iron limited samples. Since ETR_k_ does not directly include σ_PSII_, we speculate that C-assimilation will show less decoupling with this productivity metric under low iron conditions. This hypothesis remains to be tested in future studies.

### Computational considerations

Beyond the physiological and taxonomic effects described above, the computational procedures used to analyze Chl*a* fluorescence relaxation kinetics and derive turnover rates of electron transport molecules may also have a direct effect on the observed relationship between ETR_k_ and ETR_a_. We derived τ_Qa_ parameters used to calculate ETR_k_ using the FRRf fluorescence transient fitting approach, as outlined in ‘*Electron Transport Rates, ETR_PSII_′* Materials and Methods section. This approach relies on numerically fitting the rate of change in the redox state of Q_a_, driven by electron fluxes in and out of PSII reaction centers.

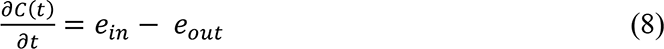

Here e_in_ is equivalent to the rate of primary photochemistry induced by excitation flashlets, and e_out_ is controlled by Q_a_ reoxidation. Within FRRf Soliense software, e_in_ is formulated as,

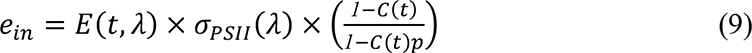

By comparison, FIRe-based analysis of fluorescence transients deviates from FRRf by including an additional term to describe reaction center closure by background actinic light (PAR) [30]. As a result, Eq. 3 is modified as:

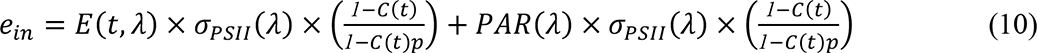

The FRRf based analysis does not include a PAR term, as it is presumed that constant background light influences the baseline of the fluorescence signal, but does not contribute to dynamic changes in fluorescence measured over the course of an ST flash [Z. Kolber, *pers. comm*.]. In this interpretation, C(t) represents the fraction of initially available reaction centers closed by excitation pulses, such that C(t=0) always equals 0.

Gorbunov and Falkowski [30] conducted a primary analysis of differences in τ_Qa_ values retrieved from FRRf and FIRe fluorescence relaxation analyses. Their results showed that when the effect of background light was explicitly included in numerical formulations, FIRe-derived τ_Qa_ values displayed strong actinic light dependencies, increasing with irradiance until plateauing around saturating irradiances (E_k_). By contrast, their FRRf-derived τ_Qa_ values varied little with actinic irradiance and resulted in a markedly shorter photosynthetic turnover time. Our own FRRf-derived τ_Qa_ values displayed a weak relationship with applied actinic irradiances (*r* = 0.17, *p* < 0.01, *n* = 203). It is possible that applying the FIRe model fit to our own data may have also yielded slower photosynthetic turnover times, and therefore lower ETR_k_ estimates, but it is unclear to what extent the alternative model may have affected our ETR_k_ results and the observed decoupling between ETR_k_ and ETR_a_.

Going forward, it will be important to separate physiological drivers of ETR_k_ and ETR_a_ decoupling from offsets resulting from the use of different mathematical approaches to data analysis. Discrepancies between ETR_k_ and ETR_a_ that cannot be explained by different derivations of *τ*_9!_ must be attributable to differences between the two ETR_PSII_ algorithms themselves. This raises the important question of which approach is most accurate, as neither has been established as a “gold standard”. Addressing this issue will require parallel independent measurements of PSII activity, such as gross oxygen evolution measurements from ^18^O experiments, as preformed previously for ETR_a_ (e.g. [49]) but not ETR_k_. Such fluorescence-independent validations of ETR_a_ and ETR_k_ are critically lacking, and will help elucidate the taxonomic and environmental influences on FRRf-based productivity measurements (Hughes et al., 2018). This, in turn, will be of significant practical utility to FRRf users seeking to derive ship-based primary productivity estimates.

### Conclusions and Future Recommendations

Fast Repetition Rate Fluorometry offers a means to rapidly assess both physiological status and photosynthetic electron transport rates of phytoplankton. The aim of this study was to evaluate an autonomous protocol for high resolution FRRf measurements of phytoplankton physiology, and to compare two alternative models for deriving primary productivity estimates from FRRf data. Our results demonstrate significant residual NQP effects after five minutes of low light acclimation, suggesting the need for extended low light acclimation periods, which would significantly decrease measurement frequency. In contrast, in-situ surface PAR had no effect on *F^′^q* /F*′*m measured under 150 μE m^-2^ s^-1^ of actinic light, suggesting phytoplankton rapidly approach steady state conditions in the presence of actinic light. Our findings also illustrated localized regions of elevated *F_v_*/*F_m_*, likely linked to local nitrogen loading by freshwater inputs and tidal mixing. Although the amplitude-based and kinetic-based derivations of photosynthetic electron transport rates were well correlated, absolute agreement between estimates appeared to be affected by phytoplankton photophysiology, with the two models diverging under nutrient-limited conditions.

As a first step, resolving discrepancies between ETR_a_ and ETR_k_ will require consensus regarding the analysis of raw fluorescence transient data. This aim is fundamental for consistent data reporting among the growing community of FRRf and FIRe users [31]. Second, ETR_a_ and ETR_k_ should be validated against fluorescence-independent measures of productivity. Considering that measurement of ETR_k_ does not require a dark acclimation step, short (∼ 5 min.) acclimation steps should be sufficient to accurately derive this term. Based on these findings, the ETR_k_ model, if validated against independent productivity metrics, may be advantageous for high resolution evaluations of in-situ photosynthetic rates under ambient light conditions.

## Acknowledgements

CTD station, surface PAR, and TSG data presented herein were collected by the Canadian research icebreaker CCGS Amundsen and made available by the Amundsen Science program, which is supported through Université Laval by the Canada Foundation for Innovation. We wish to also thank Maxim Gorbunov and Nina Schuback for valuable comments on this manuscript.

